# Single-Cell RNA-Seq Reveals Adventitial Fibroblast Alterations during Mouse Atherosclerosis

**DOI:** 10.1101/2024.10.05.616802

**Authors:** Lauren E. Fries, Allen Chung, Hyun K. Chang, Timothy L. Yuan, Robert C. Bauer

## Abstract

**Background:** Atherosclerotic cardiovascular disease (ASCVD) remains the leading cause of mortality in the western world despite the success of lipid lowering therapies, highlighting the need for novel lipid-independent therapeutic strategies. Genome-wide association studies (GWAS) have identified numerous genes associated with ASCVD that function in the vessel wall, suggesting that vascular cells mediate ASCVD, and that the genes and pathways essential for this vascular cell function may be novel therapeutic targets for the treatment of ASCVD. Furthermore, some of these implicated genes appear to function in the adventitial layer of the vasculature, suggesting these cells are able to potentiate ASCVD.

**Methods:** To investigate the role of adventitial cells in atherosclerosis, we conducted single-cell RNA sequencing (scRNA-seq) of the aortic adventitia during atherogenesis in male *Ldlr*^-/-^ mice via pools of three mice, two samples per condition. We cross-referenced the scRNA-seq data with human ASCVD GWAS to identify regulators of adventitial responses in ASCVD. These regulators were then validated *in vitro* in human adventitial fibroblasts.

**Results:** We identified four adventitial fibroblast populations, all of which displayed shifts in population size and gene expression over the course of atherogenesis. *SERPINH1*, an ASCVD-linked GWAS gene, was differentially expressed in adventitial fibroblasts during atherogenesis. Knockdown of *SERPINH1 in vitro* reduced fibroblast migration and altered subcluster marker gene expression.

**Conclusions:** These findings reveal dynamic changes in adventitial fibroblasts during atherosclerosis and suggest that reduced *SERPINH1* expression disrupts adventitial fibroblast function, contributing to ASCVD progression.

## Introduction

Atherosclerotic cardiovascular disease (ASCVD) remains the leading cause of death in the United States ^1^. The infiltration of low-density lipoprotein (LDL) particles into the vascular intima is a crucial initiating event in atherosclerosis, and therefore therapeutic strategies have traditionally focused on reducing circulating LDL levels. Despite the widespread use of LDL-lowering therapies, residual mortality from ASCVD remains high. Thus, there is a strong interest in developing novel non-LDL-lowering therapies ^2^. Unbiased genome-wide association studies (GWAS) have successfully identified novel genes and biological pathways associated with ASCVD, providing candidate novel therapeutic avenues ^3^. Notably, while GWAS have routinely identified genes involved in LDL metabolism, many genes function in pathways related to inflammation and vascular cell biology. While targeting LDL has been the gold standard in treating ASCVD, clinical trials have demonstrated that reducing inflammation is therapeutically efficacious, emphasizing the value of GWAS insights in uncovering new pathways to mitigate disease ^4^.

Many ASCVD GWAS-identified genes function in the vasculature, a finding that is perhaps unsurprising given that the vasculature is the site of atherosclerotic lesion development ^5,6^. However, the cellular and spatial complexity of the vasculature complicates our understanding of these genes’ specific roles. Arteries consist of three primary layers: the intima, which is the innermost layer and comprises predominantly a monolayer of endothelial cells; the media, the middle layer dominated by smooth muscle cells; and lastly the adventitia, the outermost layer consisting mainly of fibroblasts but also comprising multiple cell types, including smooth muscle cells, T cells, B cells, macrophages, and endothelial cells ^7^. While recent studies have explored the role of intimal and medial cells in atherosclerosis, the adventitia has received far less attention. Notably, multiple ASCVD-associated GWAS genes, including *ADAMTS7 and TCF21* ^8,9^, are expressed in the adventitia in both mouse and human atherosclerosis. Furthermore, *in vivo* mouse studies have shown that transplanted fibroblasts can migrate from the adventitia into the neointima after vascular balloon injury ^10^. These findings support a more substantial role of the adventitia in atherogenesis than previously appreciated, and suggest that understanding the role of the adventitia during atherogenesis may reveal novel therapeutic avenues.

In this study, we sought to leverage single-cell RNA sequencing (scRNA-seq) to characterize the transcriptional landscape of the vascular adventitia during atherosclerosis development in the LDL receptor knockout (*Ldlr ^-/-^*) mouse model of atherosclerosis. Our results revealed dynamic changes in adventitial fibroblasts during atherogenesis. We identified *Serpinh1*, an ASCVD GWAS gene, as differentially expressed in adventitial fibroblasts during atherogenesis. Functional studies further indicate that *SERPINH1* promotes fibroblast migration and proliferation. These findings establish a link between human ASCVD GWAS data and adventitial biology, underscoring the critical role of the adventitia in atherosclerosis and revealing a potential novel therapeutic target for ASCVD.

## Methods

### Animal studies

The Institutional Animal Care and Use Committee of Columbia University approved all animal experiments under protocol AAAZ4473. *Ldlr*^-/-^ mice (Stock No. 002207) were purchased from the Jackson Laboratory and maintained at Columbia. Mice were backcrossed for at least 10 generations as detailed by the Jackson Labs. To minimize genetic drift, inbreeding at Columbia University was restricted to no more than 10 generations. Mice were housed in an AAALAC certified vivarium in the Institute for Comparative Medicine at Columbia University, and provided the same diet and housing conditions as described here and per IACUC guidelines. Male *Ldlr*^-/^ mice, aged 8–10 weeks, were placed on a Western Diet (Research Diets, D12079Bi) for 9 or 16 weeks to induce atherosclerosis. For each experimental condition, samples were generated by pooling three mice, with two pooled samples collected per time point.

### Preparation of mouse aortic adventitia for scRNA-seq

Mice were deeply sedated with isoflurane and euthanized by cervical dislocation. Immediately following euthanasia, mice were perfused with ice-cold DPBS. The aorta was dissected from the ascending aorta to the bifurcation at the common iliac artery. The aorta was digested in 1 mg/mL of Collagenase A (Roche, 11088793001) in F12:DMEM (Cytiva, SH30023.01) containing 0.2% BSA (Sigma-Aldrich, A9647) at 37℃ to allow for the removal of the adventitia. After 5 minutes of enzymatic digestion, the aorta was moved to DPBS, and the outer adventitia was manually peeled off. The adventitia from three mice was pooled and subsequently digested in a cocktail containing 2 mg/mL Collagenase A, 0.05 mg/mL Elastase (Worthington Biochemical, LS006365), and 60 U/mL DNase I (Worthington Biochemical, LS006331) in F12:DMEM with 0.2% BSA. This digestion was performed at 37°C for 30 minutes with occasional gentle agitation.The cells were subsequently passed through a 70uM cell strainer and spun down at 300 x g for 5 minutes. The cells were resuspended in F12: DMEM containing 0.2% BSA and 1mg/ml Dnase. Cells were stained with 1:10,000 DAPI and 1:1,000 Draq5 (BioLegend, 424101) to identify live cells. Live, single cells were subsequently sorted using fluorescence-activated cell sorting (FACS). All scRNA-seq experiments were performed at the JP Sulzberger Genome Center at Columbia University. Samples were prepared using 10x Genomics Chromium Cell 3’ Reagent Kits according to the manufacturer’s instructions. A target of 5,000 cells per sample was loaded, aiming for a sequencing depth of 100 million reads per sample. Sequencing was performed on an Illumina NovaSeq 6000 platform.

### Analysis of mouse scRNA-seq data

#### Data pre-processing and quality control

Fastq files were processed with Cell Ranger 5.0.1 from 10X genomics to generate count matrices of unique molecular identifiers (UMIs). Reads were mapped to the GRCh38 human reference genome and the transcript annotated by GENCODE v32 from 10X prebuilt reference package.

We performed filtering on each sample with Seurat v5.1.0, excluding cells with < 100 features, UMI count < 500, number of genes < 200, log10GenesperUMI < 0.8, mitochondrial ratio >0.1, any genes expressed in 10 or fewer cells or with zero counts from further analyses. The samples had the following cell counts post-filtering: W0R1: 7682; W0R2: 6314; W9R1: 5063; W9R2: 9551; W16R1: 3696; W16R2: 4720. We integrated the data using SCTransform and performed integration with Harmony v1.2.1. We then formed clusters using the top 21 principal components for the shared nearest neighbor (SNN) analysis and a resolution of 0.4. To identify cell types we found conserved marker genes for the clusters with the “FindConservedMarkers” function along with utilizing previously established cell type markers from the literature. We identified six broad cell types, including B cells, T cells, fibroblasts, smooth muscle cells, macrophages, and endothelial cells **(Table S1)**. We merged all separate clusters into these six broad groups. Fibroblasts were subclustered and reprocessed to obtain 4 distinct clusters at a resolution of 0.2 after two sets of two clusters were combined due to minimal differences in conserved genes. Conserved markers were also found for these clusters **(Table S2)**.

#### Cell Type Proportions

To test for statistical significance of population size differences between clusters, we used the propeller function from the speckle v1.4.0 package, which performs a 2 way ANOVA on the cell counts **(Table S3-4)** ^11^.

#### Differential gene expression

Raw RNA counts were used to identify differentially expressed genes between time points with Nebula v1.5.3 using the standard h-likelihood method. P-values were adjusted by the Benjamini-Hochberg method **(Table S5-8, Table S9-12)** ^12^.

#### Gene set enrichment analysis

We performed Gene ontology (GO) enrichment analysis using clusterProfiler v4.12.6. We used gseGO function to perform this analysis on the differentially expressed genes between the WD time points across all clusters, as well as in fibroblasts subset specifically. We utilized only the genes commons between WD W9 and W16 and used the averaged log_2_FoldChange (log_2_FC) of the two time points. We set the number of permutations to 10000 and seed to 4444. We simplified the GO terms using a cutoff of 0.7 by the adjusted p-value. To examine the upregulated vs downregulated GO terms we used the enrichGO function, setting all genes with logFC_avg more than 0 as upregulated and those less than 0 as downregulated. We provide visualization of the results with the dotplot function.

#### Pseudotime analysis

We used monocle3 v 1.3.7 to complete pseudotime analysis of the fibroblast subcluster. Due to the stemness of multiple marker genes in the progenitor population, we set that as the root node.

### Analysis of human scRNA-seq data

#### Quality control and data pre-processing

scRNA-seq samples were obtained from the Gene Expression Omnibus (GEO, GSE159677) ^13^. All patients presented with severe carotid plaque formation, which required carotid endarterectomy. Near full thickness (except adventitia) sections of artery and plaque were recovered from three patients from the atherosclerotic core, based on the surgeon’s determination of area of largest plaque burden. Patient-matched proximal adjacent portions of carotid artery tissue were used as the control. The aggregate filtered feature-barcode matrix of the processed single-cell data was pre-processed using Seurat v5.1.0 for quality control, normalization, dimensionality reduction, and clustering. We excluded cells with < 100 features, UMI count < 500, number of genes < 200, log10GenesperUMI < 0.8, mitochondrial ratio >0.1, any genes expressed in 10 or fewer cells or with zero counts from further analyses. Samples were normalized using SCTransform in Seurat, integration was completed with HarmonyIntegration, and dimensionality reduction was performed with standard Seurat processing pipelines. We formed clusters using the top 14 principal components for the shared nearest neighbor (SNN) analysis and a resolution of 0.2. To identify cell types we used the markers the original Alsaigh et al paper listed. We identified six cell types, including smooth muscle cells, fibroblasts, endothelial cells, macrophages, B cells, and T cells.

#### Differential gene expression

Raw RNA counts were used to identify differentially expressed genes between plaque and proximal samples with Nebula v1.5.3 using the standard h-likelihood method. P-values were adjusted by the Benjamini-Hochberg method **(Table S13-14)** ^12^.

#### Locus Zoom

We used locus zoom to create the LocusZoom plot ^14^.

### Filtering CAD GWAS genes

We started off with ST10 from Aragam et al, the CAD associated GWAS genes at 1% false discovery rate (FDR) ^15^. After selecting the full list of what they deemed to be the nearest gene to the GWAS SNP, we filtered for unique genes, ending with a list of 715 genes. We then pulled the list of differentially expressed genes between WD weeks in the fibroblast cluster specifically and filtered for the list of 715 GWAS genes. We found the common genes between WD W9 and W16 and then filtered for genes with p-adjusted values at both time points less than 0.05, resulting in a list of 134 significantly differentially expressed genes. We next filtered GTEX v8 bulk RNA-seq TPM data v1.1.9 (obtained from the GTEx Portal on 12/15/2023 and dbGaP accession number phs000424.v8.p2) for genes expressed in either the coronary artery or aorta at more than 1 TPM, resulting in a list of 17115 genes. We filtered the list of CAD GWAS significantly differentially expressed genes for only those included in the GTEX set, resulting in 129 genes expressed in relevant human tissue. We then filtered for those that have more than 1.25 fold change (log_2_(0.32)) in either WD W0 vs W9 or vs W16, leaving 74 genes. We next removed from the list any genes associated with lipid traits by using a lipid associated GWAS list of 1182 genes ^16^, as we are focused on lipid-independent effects on ASCVD, leaving 65 genes. Reactome pathway analysis was completed on the list of 65 genes using enrichPathway to examine what pathways are enriched in the CAD GWAS differentially expressed genes. Using our single cell mouse data, we used the AverageExpression function from Seurat to create a matrix of average gene expression, and then created a logical matrix indicating high expression levels if the cell type has greater expression than 1 based on SCT normalization. We then filtered the matrix for those genes with high expression in fibroblast and in less than 2 other cell types. This list of 290 genes was used to filter the CAD GWAS filtered list of 65 genes to result in a final list of 6 genes.

### Cell Culture

Human aortic adventitial fibroblasts were purchased from Lonza (Lonza, CC-7014) and cultured in the Stromal Cell Growth Basal Medium containing Stromal Cell Growth Medium SingleQuots Supplements and Growth Factors (Lonza, CC-3205). Only cells between passages 3-7 were used for experiments. Cells were routinely tested for mycoplasma contamination.

### siRNA knockdown

All siRNAs were purchased from Integrated DNA Technologies (Integrated DNA Technologies, hs.Ri.SERPINH1.13). Human fibroblasts were seeded on a 12-well plate and transfection was performed using RNAiMAX (Thermo, 13778075). siRNAs were transfected at a concentration of 10nM end concentration, and knockdown or further assays were assessed 48 hours post-transfection.

### qRT-PCR

RNA from whole cell lysates was extracted using the Directzol Miniprep Kit (Zymo). The cDNA was subsequently synthesized using High-Capacity cDNA Reverse Transcription Kit (Applied Biosystems, 4368813) according to the manufacturer’s instructions. Quantitative PCR was performed using Taqman probes on the QuantStudio 7 Flex Real-Time PCR System. Analysis was performed using the ΔΔCT method with normalization against the housekeeping gene Gapdh. n=4 pooled samples of 3 wells each per condition collected after the completion of migration assay, two-tailed unpaired t-test was completed to analyze differences in gene expression between conditions. **** P<0.0001, *** P<0.001, ** P<0.01, ns P>0.05.

### Proliferation Assay

Cellular proliferation was assessed with the CellTiter 96 AQueous One Solution Cell Proliferation solution (Promega, G3582) according to manufacturers’ instructions. In brief, fibroblasts were seeded in a 96-well plate at 5,000 cells/well. The CellTiter reagent was diluted in culture media, and cells were incubated for 2 hours at 37℃ in a humidified incubator. The absorbance was subsequently measured at 490nm. n=8 biological samples per condition and a two-tailed unpaired t-test was completed to analyze differences in proliferative activity between conditions. **** P<0.0001, *** P<0.001, ** P<0.01, ns P>0.05.

### Migration assay

Cellular migration was assessed using the Culture-Insert 2 Well 24 (Ibidi, 80242). Cells were seeded at 20,000 cells/well (one half of insert) 24 hours before migration was assessed, at which point images were obtained using a Nikon Ti-S Automated Inverted Microscope at 10X magnification. Image processing was performed using ImageJ using Wound Healing Size Tool macro on manual mode and measurements are available in supplemental **Table S15** ^17^. The full set of original images is available in the supplemental images folder, n=24 biological samples (wells) per condition, completed over two separate dates, and two-tailed unpaired t-test was completed to analyze differences in migration between conditions. **** P<0.0001, *** P<0.001, ** P<0.01, ns P>0.05.

### Statistical Analysis

Values are expressed as mean±SEM. An unpaired 2-tailed Student *t* test was used to assess statistically differences between 2 groups. Values <0.05 were considered to be statistically significant. All statistical calculations were performed on GraphPad Prism.

## Data availability statement

All sequencing data will be available at a GEO accession number GSExxxx. All R scripts used for data analysis are publicly available on github at .

## Results

### ScRNA-seq of the Adventitia of Atherosclerotic *Ldlr^-/-^* Mice

To investigate the effects of atherosclerosis progression on the adventitia, *Ldlr^-/-^* mice were fed a Western Diet (WD) for 0, 9, or 16 weeks (W0, W9, W16). At each time point, the aorta was dissected, and the adventitia was mechanically isolated and dissociated for scRNA-seq **(Fig. 1A)**. Following sequencing, quality control, and UMAP clustering, we employed previously established cell lineage markers and identified six major cell types: T cells, B cells, macrophages, fibroblasts, smooth muscle cells, and endothelial cells **(Fig. 1B)**. Notably, atherogenesis caused a trend of fibroblast contraction and an expansion of immune cells **(Fig. 1C; Table S3)**. While these shifts were apparent, they were not statistically significant across the experimental timeline **(Fig. 1C; Table S3)**. Overall, fibroblasts represented the largest proportion of cells **(Fig. 1C)**.

**Fig. 1:**
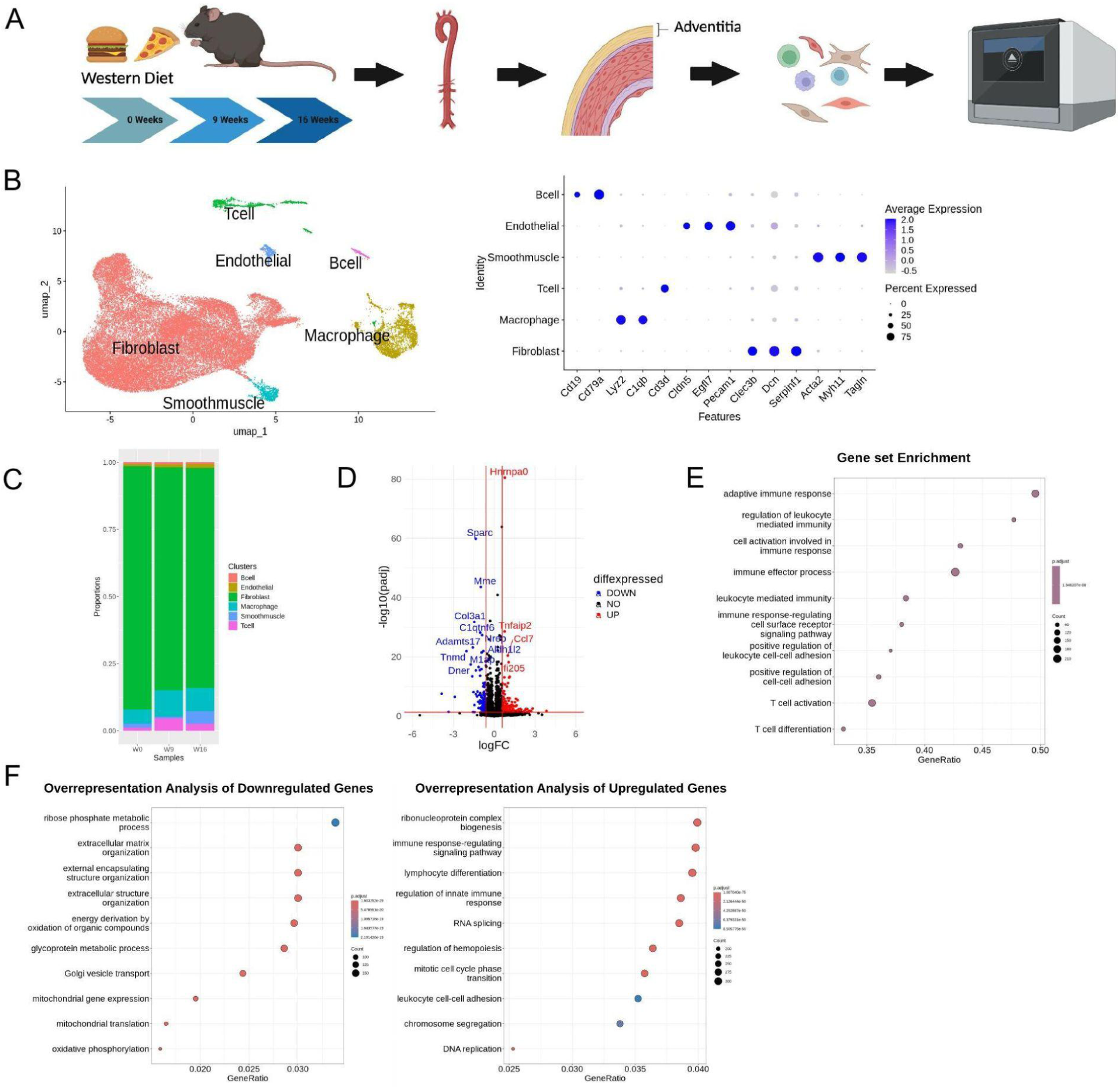
Adventitial cells during atherosclerosis progression. **(A)** Schematic of experimental design. Aortic adventia samples were collected from *Ldlr^-/-^* male mice after 0, 9, and 16 weeks (W0, W9, W16) of western diet (WD). The samples were enzymatically digested to isolate single cells for single-cell RNA-sequencing (scRNA-seq). Created with Biorender.com. **(B)** Uniform Manifold Approximation and Projections (UMAP) visualization of the filtered single cells of all three WD time points. Single-cell clustering reveals six major cell types and top marker genes to aid in cell type identification, n=2 samples per each WD time point, with 3 mice combined per sample. **(C)** A bar plot showing the proportion of each cell type across WD feeding. **(D)** Volcano plot of differentially expressed genes common between WD W0 vs W9 and W16. log2FC values are on the x-axis and the y-axis shows the difference between proportions. The left side of the x-axis represents a decrease, and the right side represents an increase in expression. **(E)** Gene set enrichment of differentially expressed genes common between WD W0 vs W9 and W16. **(F)** Overrepresentation analysis of up and down-regulated genes between WD W0 vs W9 & W16.

Examination of the differentially expressed (DE) genes between time points revealed 1,796 genes differentially expressed between week 0 and the two later time points, including multiple CAD-associated GWAS genes **(Fig. 1D; Table S6)**. Gene set enrichment analysis (GSEA) of DE genes between W0 and later time points (W9 and W16) revealed remodeling of immune and extracellular matrix (ECM)-associated processes during atherogenesis **(Fig. 1E)** ^18^. Specifically, immune-related processes were upregulated, while ECM-related processes were downregulated, aligning with the observed immune cell expansion and fibroblast contraction **(Fig. 1F)**. Collectively, our data demonstrates substantial transcriptomic remodeling of the adventitia during atherosclerosis progression. These findings highlight the adventitia as a dynamic and actively remodeling layer in atherogenesis, with significant shifts in cellular composition and molecular pathways.

### Subsetting Fibroblasts Identifies Four Distinct and Biologically Relevant Fibroblast Populations

Fibroblasts constitute the majority of adventitial cells, and prior literature has implicated adventitial fibroblasts in atherogenesis ^19^. To further investigate adventitial fibroblasts in atherosclerosis, we subclustered and re-analyzed the fibroblast population, and identified four fibroblast clusters that we have labeled ProgenitorFbs, MatrixFbs, RemodelFbs, and APCFbs **(Fig. 2A)**.

**Fig. 2:**
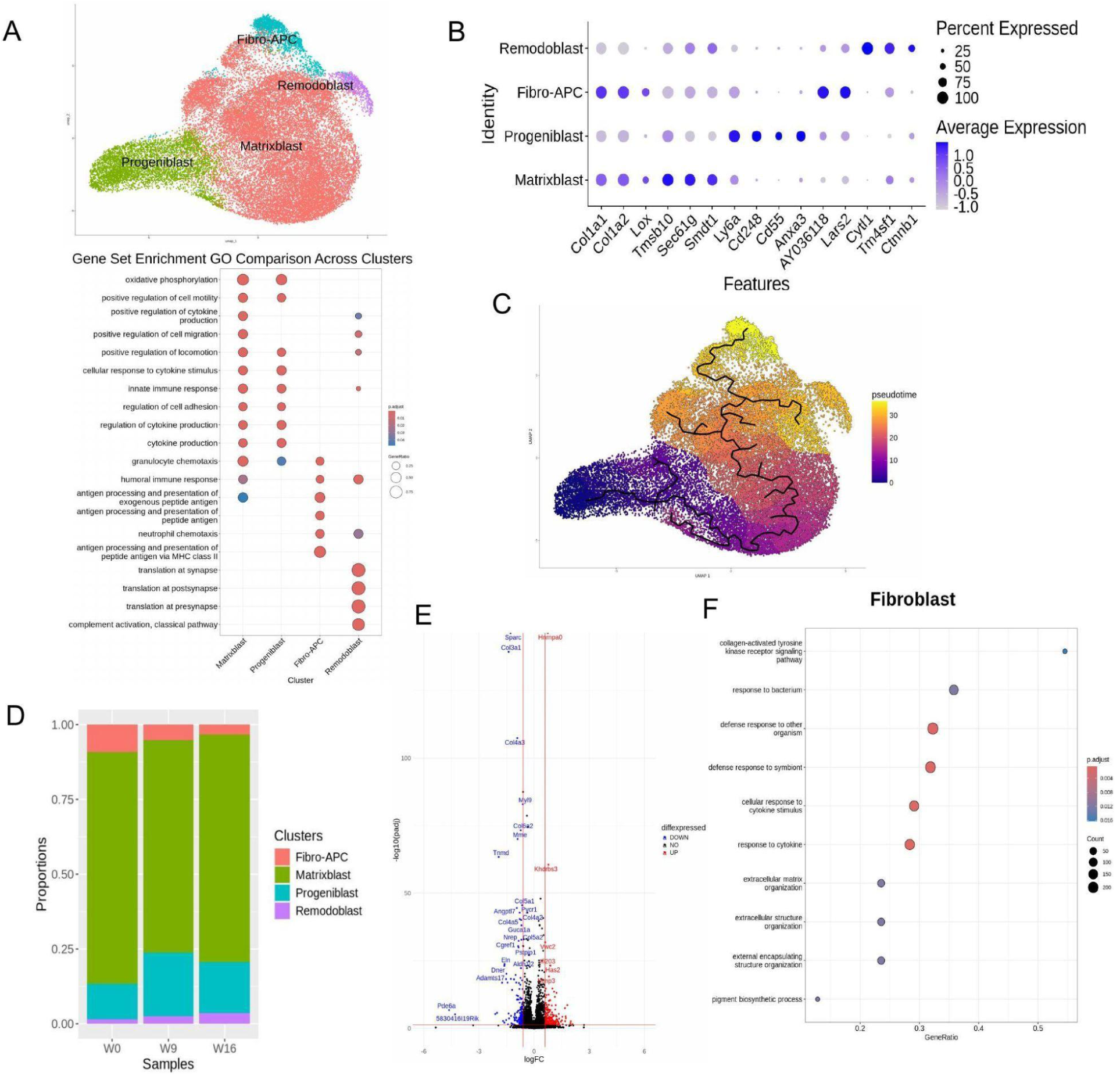
Diversity of adventitial fibroblasts and changes during atherogenesis. **(A)** Uniform Manifold Approximation and Projection (UMAP) visualization of the fibroblast subset, uncovering four populations. **(B)** A dot plot visualizing normalized expression patterns of distinguishing gene features and gene set enrichment (gseGO) analysis of conserved marker genes of each cluster. **(C)** Monocle prediction of pseudotime trajectory analysis of fibroblast subsets. **(D)** A bar graph showing the proportion differences of fibroblast subsets across WD feeding. **(E)** Top differentially expressed common genes between western diet (WD) week 0 (W0) vs. W9 and W0 vs. W16, specifically in fibroblasts. **(F)** Gene set enrichment analysis of common genes between WD W0 vs. W9 and W0 vs. W16 in fibroblasts.

MatrixFbs were characterized by high expression of *Col1a1* and *Col1a2*, markers of extracellular matrix (ECM) production, alongside *Lox*, an ECM-crosslinking enzyme ^20,21^. In addition, *Tmbsb10* and *Sec61g*, proteins involved in cytoskeleton regulation and protein synthesis/translocation, were increased, further highlighting the role of MatrixFbs in ECM production and remodeling ^22,23^. ProgenitorFbs displayed high levels of *Ly6a*, *Cd248*, and *Cd55*. *Ly6a* is a stem-like marker and, along with *Cd248,* a marker of fibroblast progenitors ^19,20,24^. *Cd55* was previously used to mark a specific fibroblast cluster described as relating to vascular development ^21^. Taken together, these markers indicate that ProgenitorFbs represent a progenitor-like fibroblast subset. RemodelFbs uniquely expressed *Cytl1*, *Tm4sf1*, and *Ctnnb1*. *Ctnnb1* is a stemness-associated gene, while *Tm4sf1* plays a role in activation and motility, and *Cytl1* is essential in ECM remodeling ^20,25,26^. These features suggest that RemodelFbs may migrate to sites requiring ECM remodeling. Finally, APCFbs, or antigen-presenting fibroblasts, express *AY036118* and *Lars2*. *AY036118* was previously labeled as a marker of an immune cell population similar to that of autoreactive memory CD8 T cells, indicating a potential immune-related role ^27^. Similarly, *Lars2* marks a subset of autoimmune B cells ^28^. In addition, GSEA of APCFbs revealed a strong enrichment for antigen presentation pathways, supporting a potential immunomodulatory role **(Fig. 2B; Table S2)**.

We next performed pseudotime analysis to identify transitions between fibroblast clusters. Pseudotime analysis revealed a transition from stem cell-like ProgenitorFbs to the more differentiated states of APCFbs and RemodelFbs **(Fig. 2C)**. Throughout atherosclerosis progression, the APCFb cluster contracted, while the RemodelFb population expanded, suggesting these subtypes potentially play a role in the cellular response to plaque formation **(Fig. 2D; Table S4)**. The population sizes of ProgenitorFbs and MatrixFbs remained unchanged **(Table S4)**. Differential gene expression analysis across clusters during WD feeding (W0, W9, and W16) identified 2,615 significantly altered genes **(Fig. 2E; Table S7-8)**. Several genes were previously linked to ASCVD, including *Apoe*, *Col4a1*, *Lpl*, and *Abca1*, but many of these genes are known to alter ASCVD through lipid-related functions. Other identified genes, including *Hnrnpa3* and *Serpinh1*, have less defined roles connecting them to ASCVD **(Table S7-8)**. In addition, GSEA of DE genes in fibroblasts highlighted substantial changes in immune and ECM-associated genes during plaque development **(Fig. 2F)**. These findings reveal the functional diversity of adventitial fibroblast populations and underscore their cellular alterations during atherosclerosis progression.

### Identification of *SERPINH1* as an Adventitial Fibroblast Gene of Interest

Given that ASCVD GWAS genes have been previously shown to be expressed in adventitial cells during atherosclerosis, we aimed to identify ASCVD GWAS-identified genes that were differentially expressed in our adventitial fibroblast dataset. To achieve this, we built a pipeline beginning with coronary artery disease (CAD) associated GWAS genes ^15^ and integrated this list with adventitial fibroblast-specific differentially expressed genes from our scRNA-seq data **(Fig. 3A)**.

**Fig. 3:**
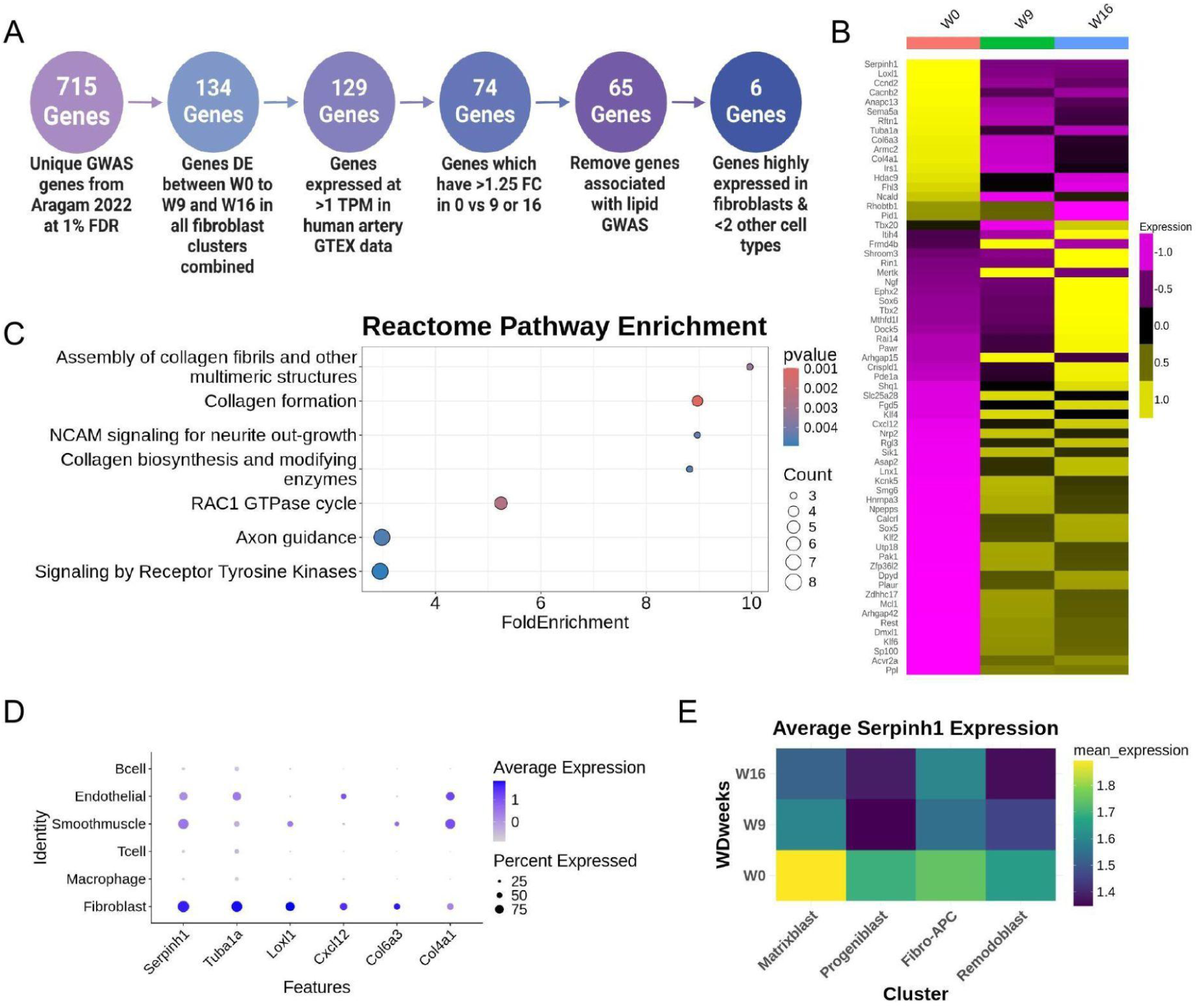
CAD GWAS gene cross analysis with expression changes during atherosclerosis in mice. **(A)** Computational pipeline for cross-referencing CAD GWAS hits and differentially expressed adventitial fibroblast scRNA-seq identified genes. **(B)** Heatmap of expression of the 65 differentially expressed CAD GWAS genes in the adventitia across WD feeding after removing lipid GWAS associated genes. **(C)** Gene set enrichment analysis of the 65 cross-referenced genes between CAD GWAS and scRNA-seq differentially expressed in the adventitia after removing lipid GWAS associated genes. **(D)** Dot plot of the expression pattern of the six identified adventitial fibroblast CAD GWAS genes of interest. **(E)** Average expression of *Serpinh1* within fibroblast subsets across WD feeding.

The published GWAS study identified 715 unique genes at a 1% false discovery rate. We cross-referenced this list with genes significantly DE across time points in fibroblasts from our adventitia dataset, narrowing the list to 134 overlapping genes. These were further filtered by removing genes with less than 1 transcript per million (TPM) expression in the human aortic and coronary artery Genotype-Tissue Expression (GTEx) project RNA-seq data set, leaving 129 genes. To focus on genes with substantial expression changes during atherosclerosis progression, we retained only those with a fold change >1.25 in W0 vs either W9 or W16, reducing the list to 74 genes. Next, we removed any genes previously associated by GWAS with plasma lipid levels, leaving 65 genes ^16^. Of these 65 genes, approximately one-third were downregulated from WD W0 to W9 and W16, while the other two-thirds were upregulated during the progression of atherosclerosis **(Fig. 3B)**. Gene ontology (GO) enrichment was performed on this list of 65 cross-referenced genes, revealing that these genes primarily mediate ECM remodeling, highlighting the role of the adventitia in this process **(Fig. 3C)**. Finally, to identify fibroblast-specific candidates, we only retained genes expressed at >1 corrected UMI in fibroblasts and in fewer than two other cell types, resulting in a final list of six genes we consider to be fibroblast-enriched: *Serpinh1, Tuba1a, Loxl1, Cxcl12, Col6a3,* and *Col4a1* **(Fig. 3D)**.

All 6 of these genes are highly expressed in all identified fibroblast subtypes, although *Cxcl2*, *Col6a3*, and *Col4a1* are only expressed in ∼50% of fibroblasts, making them potentially less relevant. *Col4a1* encodes a type IV collagen ɑ protein, and genetic variants in it are associated with increased risk of myocardial infarction, atherosclerotic plaque instability, and vascular cell survival ^29^. *Col6a3* encodes a chain of type VI collagen, which binds extracellular matrix proteins and can contribute to thicker plaque caps and therefore more stable plaque phenotypes ^30^. *Cxcl12* encodes a member of the intercrine family, playing a role in numerous cellular functions, including inflammation response. Statins are known to reduce CXCL12 levels and patients with higher CXCL12 levels plus atherosclerosis are at an increased risk for developing thrombosis ^31^. *Loxl1* encodes a member of the lysyl oxidase family of proteins and is essential for the formation of cross-links in collagen and elastin, and *Loxl1* smooth muscle cell conditional knockout mice show protections against atherosclerotic plaques and calcification ^32^. *Tuba1a* helps form microtubules and is essential for cell division and movement, though no connection to ASCVD has been found ^33^. Finally, *Serpinh1* encodes a member of the serpine superfamily of serine proteinase inhibitors and is crucial in controlling collagen synthesis ^34^.

*Serpinh1* stood out as a gene of interest due to its high expression levels across all fibroblast subtypes. Notably, *Serpinh1* expression is statistically reduced across all fibroblasts during atherosclerosis progression, though MatrixFbs and APCFbs show higher expression levels compared to the other two fibroblast populations **(Fig. 3E; Table S7-8)**. While smooth muscle cells and endothelial cells display some *Serpinh1* expression, their expression does not change during atherosclerosis, as determined by DE analysis **(Table S9-12)**. These findings indicate that *Serpinh1* is an adventitial fibroblast gene whose expression is altered and downregulated during atherosclerosis. To determine the clinical relevance of our findings, we sought to corroborate these results in humans.

### *SERPINH1* in Human Atherosclerosis

We next interrogated available human data pertaining to *SERPINH1* in ASCVD. The lead single nucleotide polymorphism (SNP) from the coronary artery disease (CAD) GWAS, rs584961, is a synonymous coding variant in exon 2 of SERPINH1, which is consistent with SERPINH1 being the causal gene at this GWAS locus ^14,15^ **(Fig. 4A)**. Furthermore, expression quantitative trait loci (eQTL) analysis in cultured fibroblasts indicates that the risk allele (G) is associated with decreased *SERPINH1* expression **(Fig. 4B)**, reinforcing the potential anti-atherogenic role of *SERPINH1* ^15^. These observations are consistent with our observation in mice that fibroblast *Serpinh1* expression decreases over the course of atherogenesis, suggesting that the GWAS association between ASCVD and *SERPINH1* may be, in part, explained by its function in fibroblasts.

**Fig. 4:**
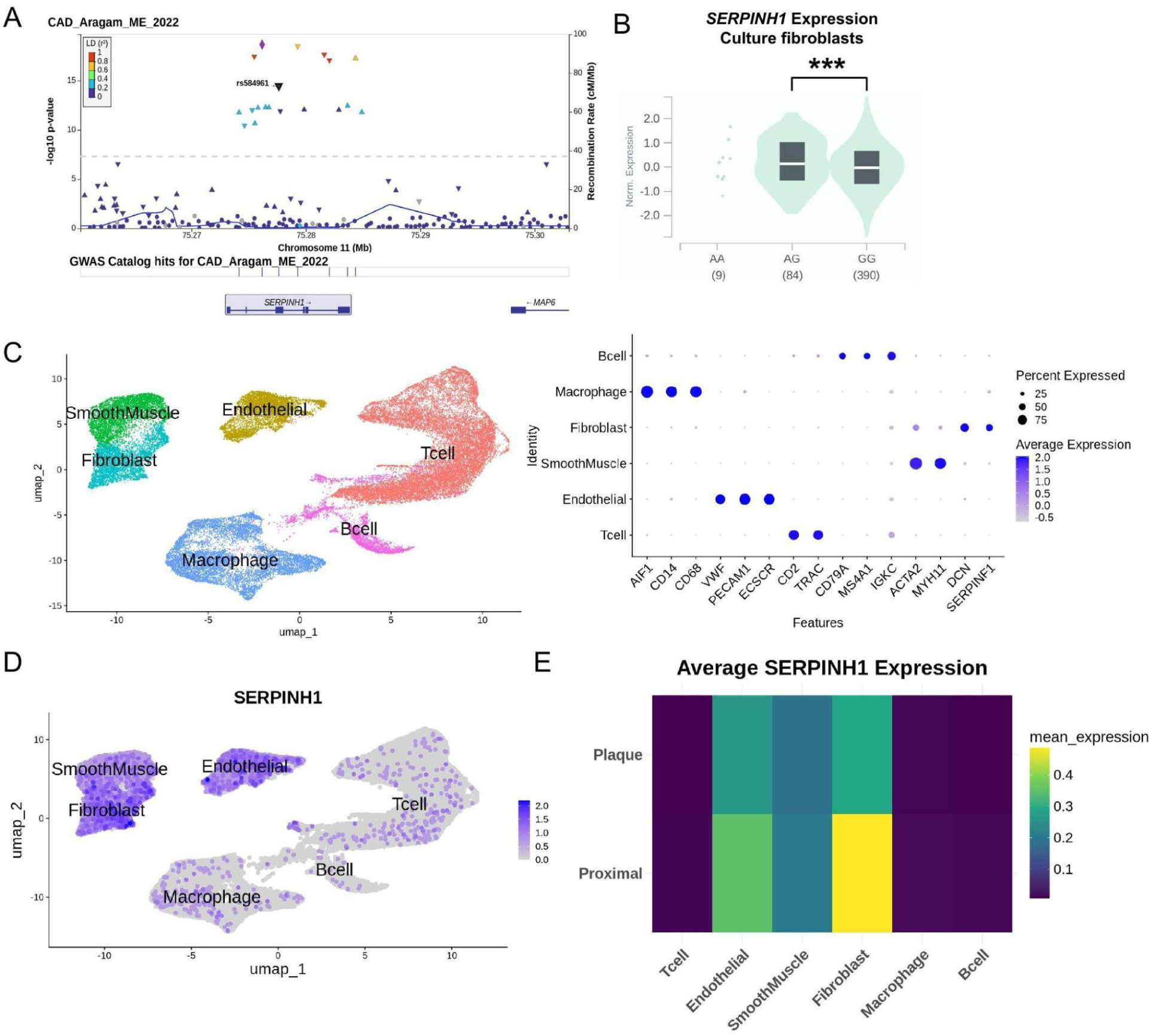
Examination of *SERPINH1* in humans during atherosclerosis. **(A)** Locus zoom plot of *SERPINH1*. **(B)** eQTL of *SERPINH1* SNP derived from GTEx (pvalue=9.84e-18). **(C)** Basic clustering identities of the joint analysis between proximal and plaque samples of human carotid arteries. **(D)** Expression of *SERPINH1* across different cell types. **(E)** *SERPINH1* average expression between the two plaque states across cell types.

Next, we set out to determine if the cell type expression patterns we observed in our mouse scRNA-seq data were reflected in human disease. To address this, we reanalyzed a previously published scRNA-seq dataset from three patients undergoing carotid endarterectomy ^13^. The study dataset included two sample types: one labeled as proximal and obtained from a full artery thickness, excluding the adventitia, while the second sample was denoted as plaque and was procured from the atherosclerotic core. We performed clustering analysis in this data and identified 6 populations: smooth muscle cells, fibroblasts, endothelial cells, macrophages, T cells, and B cells **(Fig. 4C)**.

Consistent with our observations in the *Ldlr^-/-^* mouse model of atherosclerosis, *SERPINH1* expression was predominately in fibroblasts **(Fig. 4D)**. Notably, a down-regulation of *SERPINH1* expression was observed between “proximal” healthy tissue and “plaque” data, most prominently noted in the fibroblast subset **(Fig. 4E, Table S14)**. Overall, the reduced expression of *Serpinh1* during murine atherosclerosis mirrors the reduction in *SERPINH1* expression seen in human plaques, consistent with an anti-atherogenic role for fibroblast *SERPINH1*.

### *SERPINH1* Knockdown Decreases Fibroblast Migration and Proliferation

Finally, to explore the role of *SERPINH1* in regulating adventitial fibroblast function, we performed siRNA-mediated knockdown of *SERPINH1* in primary human aortic adventitial fibroblasts (hAoFs) **(Fig. 5A)**. Knockdown of *SERPINH1* resulted in a 20% reduction in hAoFs migration as assessed by scratch assay over an 8 hour period compared to cells receiving a non-targeting control siRNA **(Fig. 5B-C, Table S15)**. Conversely, there was no change in hAoF proliferation with gene knockdown, consistent with any changes in migration being due to movement of the cells and not proliferative capacity **(Fig. 5D)**. Lastly, we evaluated the impact of *SERPINH1* knockdown on the expression of fibroblast subcluster markers. We found that *SERPINH1* knockdown causes increases in both *ANXA3* and *LOX*, markers of MatrixFb and ProgenitorFb, respectively **(Fig. 5E)**. Collectively, these results suggest that *SERPINH1* knockdown not only impairs fibroblast migration but also alters the molecular profile of hAoFs, shifting their marker gene expression.

**Fig. 5:**
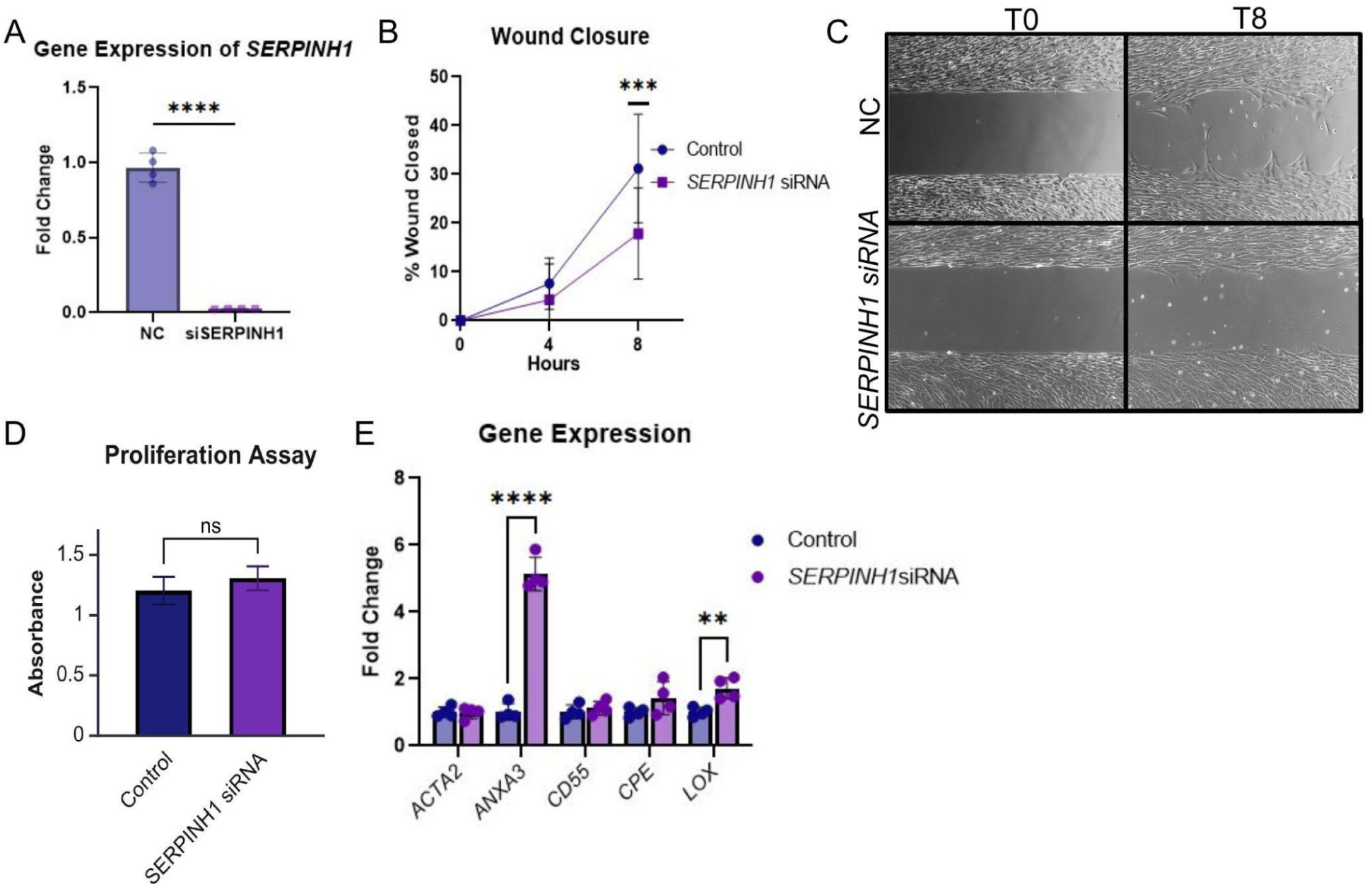
Knockdown of *SERPINH1* in human adventitial fibroblasts. **(A)** Confirmation of *SERPINH1* siRNA knockdown, n = 3 wells aggregated per sample collected after completion of migration assay. **(B)** Quantification of migration assay, n = 24 wells per condition completed on 2 separate days. **(C)** Representative image of migration assay. (D) MTT proliferation assay, n=8 wells. **(E)** Gene expression via qPCR of *ACTA2*, *ANXA3*, *CD55*, *CPE*, and *LOX,* n = 3 wells aggregated per sample collected after completion of migration assay. Normalization was performed relative to *GAPDH*. **** P<0.0001, *** P<0.001, ** P<0.01, ns P>0.05 by two-tailed unpaired t-test. Error bars show standard deviation.

## Discussion

Here, we present the first comprehensive single-cell atlas of adventitial cells through the progression of atherogenesis in hyperlipidemic *Ldlr^-/-^* mice, expanding on prior studies that focused on the *Apoe^-/-^* background or the adventitial non-leukocyte populations within *Ldlr^-/-^* mice at static time points ^21,24^. Our findings reveal that fibroblasts dominate the cellular landscape of the adventitia and undergo modest contraction in population size during atherogenesis, while adventitial immune cells exhibit a marked expansion. These cellular changes are accompanied by alterations in ECM gene expression, primarily in fibroblasts. Notably, we identify four previously uncharacterized fibroblast subclusters, including one that contracts significantly and another that expands during disease progression. Finally, we highlight *SERPINH1*, an ASCVD GWAS gene, as a likely regulatory of adventitial fibroblast function during atherosclerosis.

The role of adventitial fibroblasts in atherogenesis remains largely unexplored, with functional subsets poorly defined. While atherosclerosis has typically been viewed as an inside-out inflammatory process, emerging evidence suggests an alternative outside-in model, where inflammation originates in the adventitia and propagates to the intima ^35^. This paradigm is supported by studies showing that adventitial fibroblasts migrate to the intima following vascular injury such as balloon induced injury ^10,36^. Furthermore, multiple ASCVD GWAS-identified genes demonstrate adventitial expression in models of atherosclerosis and vascular injury ^8,9^. Understanding the role of adventitial fibroblasts and their subtypes in atherosclerosis is critical, as these cells are likely key contributors to plaque development and disease progression.

In our study, we highlight the diversity of adventitial fibroblasts throughout atherogenesis in mice. We identified four distinct adventitial fibroblast subpopulations: ProgenitorFbs, MatrixFbs, RemodelFbs, and APCFbs. Using gene expression profiling, gene set enrichment analysis (GSEA), and pseudotime analysis, we propose specific roles for each subset. ProgenitorFbs appear to represent a more transitional state, consistent with previous descriptions of fibroblast progenitors ^19,20,24^. MatrixFbs may play an important role in producing and modifying ECM proteins ^20,22,23^. APCFbs exhibit characteristics of antigen-presenting cells, implicating them in immune surveillance or modulation ^27,28^. Notably, we observed a significant contraction of the APCFb cluster over the course of atherosclerosis, suggesting a potential link between the loss of this population and disease advancement. The RemodelFb cluster likely migrates to sites of vascular injury to facilitate ECM remodeling, which is reflected in their marked expansion during atherogenesis ^20,25,26^.

While our fibroblast clusters partially align with those previously described in the literature, they do not directly correspond to previously established categories. This discrepancy likely arises from differences in analytical resolution, where our high-resolution approach using only the adventitia enabled the identification of finer subpopulation distinctions. For example, several research groups have published fibroblast atlases in both mice and humans, but as they are across different tissues, fibroblast populations are generally categorized based on the organ that the dataset is derived from ^37,38^. Moreover, prior studies have provided limited functional characterization of fibroblast subtypes, underscoring the novelty and significance of our findings.

In comparison to previously published subsets, our study demonstrates alignment with several fibroblast subclusters while also highlighting some differences. For instance, ProgenitorFbs in our dataset express *Cd55* while MatrixFbs and APCFbs expressed *Lox*, both found to be identifying markers of specific fibroblast trajectories in prior studies ^21^. Another group identified *Comp* as a marker for matrifibrocytes (or reactive fibroblasts), yet in our data, this marker is spread across all populations at low levels ^20^. Similarly, while Col4a1 and Col4a2 were proposed as markers of quiescent fibroblasts, these genes also exhibited widespread expression across all of our populations. Two groups proposed *Lrrc15*, *Mmp1*, and *Acta2* as markers for myofibroblasts, yet these genes were either not present or at very low levels without clustering in specific subsets in our dataset ^19,39^. *Gli1*, *Sca1*, and *Pdgfrb* were used as markers for fibroblasts that migrate into lesions. However, only *Pdgfrb* showed relevance in our dataset as a marker for RemodelFbs, while the other genes were expressed at very low levels or not at all ^19^. Furthermore, *Clec3*, *Dcn*, *Gsn*, and *Lum* were previously identified as markers for a specific fibroblast subcluster in the mouse aorta; however, these genes exhibited near-universal expression across all clusters in our study ^40^. Lastly, our dataset did have high universal expression of *Dpep1*, a gene used to mark adventitial specific fibroblasts^21^. These conflicting findings underscore the need for deeper sequencing in future studies, as our data set was able to delineate adventitial fibroblast subtypes and their distinct functions during atherogenesis more thoroughly and accurately than previous studies.

In this study, we cross-referenced our murine scRNA-seq with published human ASCVD GWAS datasets to identify genes with translational potential. This analysis identified *SERPINH1* as a DE gene in adventitial fibroblasts during atherosclerosis that has been linked to ASCVD in humans. *SERPINH1* encodes HSP47, a collagen-specific chaperone essential for proper collagen folding ^41^. Reduced HSP47 levels are associated with impaired cell migration and proliferation, leading to defective scar formation ^42,43^. GWAS and eQTL analysis suggests that lower *SERPINH1* expression correlates with increased CAD risk **(Fig. 4B)**. Conversely, patients with asymptomatic atherosclerosis exhibited increased expression of *SERPINH1*, highlighting a potential role for *SERPINH1* in plaque stability ^44^. In our mouse scRNA-seq, *Serpinh1* expression in adventitial fibroblasts decreases as atherosclerosis progresses. Similarly, human scRNA-seq analyses showed lower SERPINH1 levels in plaque fibroblasts compared to proximal samples, although a limitation of this dataset is that the adventitia layer was excluded from these samples. Finally, we found *in vitro* knockdown of *SERPINH1* in human adventitial fibroblasts reduced fibroblast migration and altered expression of fibroblast subcluster marker genes (*ANXA3*, *LOX*). Collectively, these findings suggest that reduced *SERPINH1* expression during atherosclerosis alters adventitial fibroblast function and identity, and may play a role in the progression of atherosclerosis.

In conclusion, our scRNA-seq analysis highlights the biological and functional diversity of adventitial fibroblasts during atherosclerosis. Our data set reveals novel functional fibroblast subtypes due to our higher resolution dataset focused solely on the adventitia during atherogenesis, of which no prior datasets exist. We highlight *SERPINH1* as an ASCVD GWAS gene with reduced expression in both mouse and human atherosclerosis, suggesting an anti-atherogenic role. *In vitro* studies showed that knockdown of *SERPINH1* in cultured human adventitial fibroblasts impairs migration and alters expression of fibroblast subcluster markers, suggesting that *SERPINH1* plays a critical role in fibroblast function and identity. Further studies in rodent models are needed to explore the role of *SERPINH1* and other CAD GWAS genes in adventitial cells during atherosclerosis. These findings underscore the emerging significance of adventitial fibroblasts and the critical role of *SERPINH1* in atherosclerosis.

## Non-standard Abbreviations and Acronyms

ASCVD: Atherosclerotic cardiovascular disease
LDL: Low-density lipoprotein
GWAS: Genome-wide association study
scRNA-seq: Single cell RNA-sequencing
*Ldlr*: Low-density lipoprotein receptor
WD: Western diet
W0 / W9 / W16: Week 0 / 9 / 16
GSEA: Gene set enrichment analysis
ECM: Extracellular matrix
DE: Differentially expressed
TPM: Transcript per million
GTEx: Genotype-Tissue Expression
GO: Gene ontology
SNP: Single nucleotide polymorphism
eQTL: Expression quantitative trait loci
CAD: Coronary artery disease
hAoF: Human aortic adventitial fibroblast

## Acknowledgements

The Genotype-Tissue Expression (GTEx) Project was supported by the Common Fund of the Office of the Director of the National Institutes of Health, and by NCI, NHGRI, NHLBI, NIDA, NIMH, and NINDS.

## Supplemental Material

Supplemental Methods Supplemental Table S1-S15 Supplemental Images Major Resources Table

## Author Contributions

LEF completed all analyses of data and *in vitro* work. AC, HKC acquired the single RNA-sequencing mouse data. AC, TY conducted experiments and acquired data. RCB oversaw the design, execution, management, and funding of all studies. LEF, AC, and RCB wrote the manuscript, which all authors approved before submission. RCB had full access to all the data in this study and takes responsibility for its integrity and the data analysis.

## Highlights

- We found significant biological and function diversity in adventitial fibroblasts during atherosclerosis progression.
- *SERPINH1* is an ASCVD GWAS gene with reduced expression in both mouse and human atherosclerosis, suggestive of an anti-atherogenic effect.
- *In vitro* knockdown of *SERPINH1* impairs human aortic adventitial fibroblast migration and markers of specific fibroblast populations, suggesting a critical role of the gene in fibroblast function and identity.

